# Targeted mRNA restoration of ciliary function in DNAI1-related primary ciliary dyskinesia: ex vivo rescue in patient-derived nasal spheroids – A pilot study

**DOI:** 10.64898/2026.05.26.727937

**Authors:** Caroline Marie Torp Nygaard, Cristian Herrera-Cid, Laura Nedergaard, Stine Green Johansen, John G Matthews, Jessica A Couch, Tavs Qvist, Kim G Nielsen, Søren Tvorup Christensen, June Kehlet Marthin

## Abstract

**Rationale:** Primary Ciliary Dyskinesia (PCD) is a genetic disorder characterized by impaired ciliary function, defective mucociliary clearance, and progressive lung disease. Pathogenic variants in the *DNAI1* gene are a well-known cause of PCD. Currently, no approved therapies address the underlying genetic defect. RCT1100 is an inhaled mRNA therapy encoding DNAI1 currently under clinical development. This study evaluates the functional effects of RCT1100 using a fast three-dimensional explant spheroid (3DE-S) model consisting of apical-out undifferentiated nasal epithelial cells derived from patients with DNAI1 PCD.

**Methods:** 3DE-S were generated from nasal brushings of five patients with confirmed biallelic *DNAI1* variants. RCT1100 was administered from day 5 directly to culture wells three times weekly for two weeks. Spheroid motility was assessed throughout treatment by quantifying the proportion of moving spheroid rolling and their movement velocity. Following six doses, spheroids were harvested for high-speed video microscopy for assessment of ciliary beat frequency.

**Results:** Evaluable data were obtained from three of five patient samples; two samples were excluded due to contamination. After six doses of RCT1100, ciliary beat frequency increased from a baseline range of 2.8-3.5 Hz to 6.7-6.8 Hz post-harvesting. Mean spheroid movement velocity increased from 0.11 µm/sec to 3.87 µm/sec following dosing with 10 µg/mL RCT1100, with more than 80% of spheroids exhibiting coordinated rolling motion pattern.

**Conclusion:** The 3DE-S is a robust platform for evaluating targeted therapies. RCT1100 significantly restored ciliary function, supporting its therapeutic potential and highlighting the utility of spheroid-based systems for precision medicine approaches in DNAI1 PCD.

## Background

Primary ciliary dyskinesia (PCD) is a genetically heterogenous disorder caused by pathogenic variants in genes essential for the structure and function of motile cilia. Inheritance is most commonly autosomal recessive, although autosomal dominant and X-linked forms have been described. Impaired ciliary motility affects multiple organ systems, with the most significant clinical burden occurring in the respiratory tract, where defective mucociliary clearance leads to chronic upper and lower airway infections. Over time, this results in progressive lung disease and declining pulmonary function.

Biallelic pathogenic variants in Dynein Axonemal Intermediate Chain 1 (*DNAI1)* represent a well-established cause of PCD)^1^. *DNAI1* encodes a 699 amino acid component of the intermediate chain of the outer dynein arm (ODA) and follows an autosomal recessive inheritance pattern. Diagnostic features of DNAI1 PCD typically include absent ODA on transmission electron microscopy (TEM) and predominantly immotile cilia on high-speed video microscopy (HSVM)^1^. Current management strategies are limited to symptomatic treatments, including airway clearance and antibiotic therapy^2^, and no approved therapies target the underlying genetic defects.

RCT1100 is an inhaled mRNA therapy encoding DNAI1, investigated in a Phase 1b clinical trial. The therapy utilizes a selective-organ-targeting (SORT) lipid nanoparticle (LNP) platform optimized for aerosol delivery to the epithelium of the lower airways via oral inhalation. Preclinical studies of RCT1100 components have demonstrated restoration of ciliary activity in air-liquid interface (ALI) cultures^3^.

In this pilot study, we applied a three-dimensional explant (3DE) spheroid method, previously established by our group^4^, as a rapid and cost-effective platform to assess ciliary motility and functional response to targeted dosing in patient-derived samples.

## Methods

### Design

This was an exploratory Single Center Pilot Study within the RCT1100-103 trial (EU Clinical Trials Register (EUCTR): 2023-510521-14-00, ClinicalTrials.gov ID: NCT06633757). the study was conducted in a collaboration between National PCD Centre, Copenhagen University Hospital, Rigshospitalet, DK, The Cilia Group, Section for Cell Biology and Physiology, University of Copenhagen, DK and ReCode Therapeutics, CA, USA.

### Participants

Participants were recruited at the University Hospital of Copenhagen between January and February 2025. Eligible participants were adults with PCD aged 18–75 years with genetically confirmed biallelic pathogenic variants in the *DNAI1* gene. Individuals with comorbidities deemed by the primary physician likely to interfere with trial participation or outcomes were excluded. Only patients included in the RCT1100-103 trial were included in the study.

### Patient sampling and nasal brush biopsy

A nasal brush was inserted into the upper third of the nasal cavity of one nostril in the patients’ nose and gently rotated for 5-10 seconds to collect ciliated epithelial cells. Immediately after nasal brushing, the cells were transferred from the brush to prewarmed medium (37 °C) consisting of DMEM/Ham’s F-12 (1:1) supplemented with a serum substitute (1% Ultroser G; IBF Biotechnics, Savage, MD, USA), antibiotics (penicillin 10^5^ U L^−1^ and streptomycin 100 mg L^−1^), and 2 mM L-glutamine.

Nasal brush biopsy samples for HSVM obtained from patients at screening for the RCT1100-103 trial were exclusively evaluated for ciliary beat pattern (CBP) and ciliary beat frequency (CBF) and served as baseline non-dosed material. A second nasal sampling for Three-dimensional Explant Spheroid formation and investigation was performed at a later occasion.

### Three-dimensional Explant Spheroid formation

In the laboratory, the collected nasal epithelial cells were gently resuspended using a pipette and distributed together with the medium into four wells of an uncoated 24-well plate (1 mL in each). The plate was then placed in a humidified CO2 incubator (5% CO2, 37 °C). To prevent attachment of the cells to the well bottom, the plate was gently tapped every 15 minutes during the first four hours of incubation and then left overnight in the CO2 incubator for formation of 3-Dimensional explant spheroids with apical out oriented cilia. Culture medium was replaced daily for the first two days and subsequently three times per week. Media exchange was performed by letting the cellular material sediment for 15 minutes in a test tube and carefully replacing the supernatant^4^.

### Dosing of spheroids

RCT1100 was pipetted into suspension at final concentrations of 1.25, 2.5, 5, 10, and 20 µg/mL of mRNA. The first dose was applied on day 5 ± 1 after spheroid formation; in samples with excessive mucus, treatment initiation was delayed by one additional day to improve culture quality by an additional change of media. Spheroids were treated three times per week for a total of six doses. For each RCT1100 dosing, spheroids were temporarily transferred to 15 mL Falcon tubes and allowed to sediment for 15 min, after which the supernatant was removed and discarded. During sedimentation, a prewarmed RCT1100-containing medium, in concentrations as specified above, was added to the wells in two aliquots to prevent drying. Spheroids were then returned to their corresponding wells and incubated with RCT1100 for 5 h at 37°C in 5% CO_2_. Following incubation, spheroids from all wells, including untreated controls, were again transferred to Falcon tubes, allowed to sediment, and the supernatant was removed prior to returning the spheroids to fresh medium. Cultures were maintained at 37 °C until the subsequent treatment 48 h later. This procedure was repeated until each well had been dosed for a total of six times over a two-week period.

### Imaging

#### HSVM

HSVM was conducted on all available samples in accordance with standardized protocols^5^. Mounted-slide high speed video analysis was used to assess ciliary beat pattern (CBP) and ciliary beat frequency (CBF) pre- and post-treatment as routinely used for diagnostic purposes at the PCD Center, Cilia Lab, Department of Paediatrics and Adolescent Medicine, Copenhagen University Hospital, Rigshospitalet. Videos were recorded at 250 frames pr second using a Leica DM4 B microscope with at DMK 33UX287 camera. Spheroids were placed between an object glass and a cover glass leaving sufficient room for cilia movement. CBP was assessed by an experienced observer using recognised nomenclature^5^. CBF was recorded in Hz using the RHcilieCore program (unpublished internal software).

#### 3DE motility

Overall spheroid motility was assessed by live cell imaging at three time points, in pre-treated cells and after the fourth and sixth dose. Imaging was performed using an IX83 Inverted Microscope equipped with a Yokogawa CSU-W1 spinning disk confocal scanner under controlled atmosphere (5% CO2 and 37°C). Spheroids were recorded for 3 minutes, using 10x phase contrast magnification and a time step of 0.6 seconds between each frame. Analysis of the spheroid motility was performed in ImageJ software v1.54p, and video was exported at 25 frames per second.

To measure translational speed, we used the TrackMate 7 plugin in ImageJ (PMID: 35654950). Each spheroid was treated as an individual particle, and its full trajectory was tracked to calculate its speed over the entire path. The mean speed of all spheroids was then computed and defined as the translational speed for each patient

### Statistical analysis

All data were summarized using descriptive statistics. Continuous variables are presented as means ± standard error of mean (SEM). Data was analysed by One-Way ANOVA. Significant differences were calculated with a P-value lower than 0.05.

## Results

Five patients aged 20-41 years were included in the study (2 females, 3 males). Samples from one patient were excluded due to contamination; as samples failed to generate viable spheroids, likely due to a concurrent upper respiratory tract infection at the time of sampling. Due to technical difficulties samples from another patient did not yield videos for HSVM. Consequently, four patient-derived samples were available for live cell imaging and three for HSVM analysis (Table 1). In all evaluable samples, the 3DE spheroids with apical out orientated cilia formed within 4-24 hours of initial incubation.

**Table 1.** Basic data.

| Patient | Sex | Age | DNAI1 variant | FEV1*<br>(Percent Predicted) | TEM*<br>(ODA / IDA) | NasalNO*<br>(ppb) | HSVM | 1-hour MCC*<br>(%) |
| --- | --- | --- | --- | --- | --- | --- | --- | --- |
| 1** | F | 41 | c.48+2dup<br>AND<br>c.1019+5G>C | 71% | NA | 68 | Decreased,<br>asynchronous | 5 |
| 2** | M | 31 | c.565_566del<br>AND<br>c.501+85G>C | 70% | 1.1 /<br>2.1 | 42 | Immotile | 1 |
| 3** | F | 23 | Bi-allelic<br>splicing<br>mutation<br>(skipping of<br>exon 6)<br>frameshift | 77% | 0.4 /<br>2.7 | 18 | Immotile | 0.7 |
| 4 | M | 20 | c.48+2dup | 95% | NA | 200 | Immotile | 3 |

\*Before entering the RCT1100 clinical trial
\*\*Samples used for HSVM analysis

### RTC110 enhances 3DE spheroid motility

Live-cell imaging was used to evaluate spheroid motility under different treatment conditions. Unlike previous methods for evaluating apical-out airway organoids (AOAO) using Matrigel-embedded spheroids^6^, we employed a “free-field” approach, allowing unrestricted spheroid movement driven by apical out cilia within the culture media.

Spheroid movement was classified as 1) non-motile, with no or minimal displacement (Figure 1A, Supplementary Video 1+2); 2) rotational/tilting motion defined as partial or complete rotation around the spheroid axis (Figure 1A, Supplementary Video 3); and 3) rolling, characterized by translational movement across the culture surface driven by coordinated ciliary activity (Figure 1B, Supplementary Video 4). After four doses, RTC1100 treatment induced a reduction in the proportion of immotile spheroids to 13.0%, 43.7% and 7.7% at 5, 10 and 20 µg/mL of RTC1100, respectively. In contrast, 92.9% of untreated spheroids were non-motile., The reduction in the non-motile spheroid fraction was translated to a significant increase in tilting/rotational movement, observed in approximately 70.0% and 64.5% of spheroids treated with 5 and 20 µg/mL of RTC1100, respectively (Figure 1B). At sixth dose, the proportion of non-motile spheroids decreased from 67.9% in untreated cultures to 19.6%, 3.3% and 11.7% at 5, 10 and 20 µg/mL, respectively. Correspondingly, rolling motion increased significantly, particularly at 10 and 20 µg/mL, where 63.8% and 59.5% of spheroids exhibited a rolling motion compared with 3.6% of untreated controls (Figure 1D). Although the 5 µg/mL dose showed early improvement after four doses, this effect was not sustained at the six-dose time point (Figure 1D).

**Figure 1.**
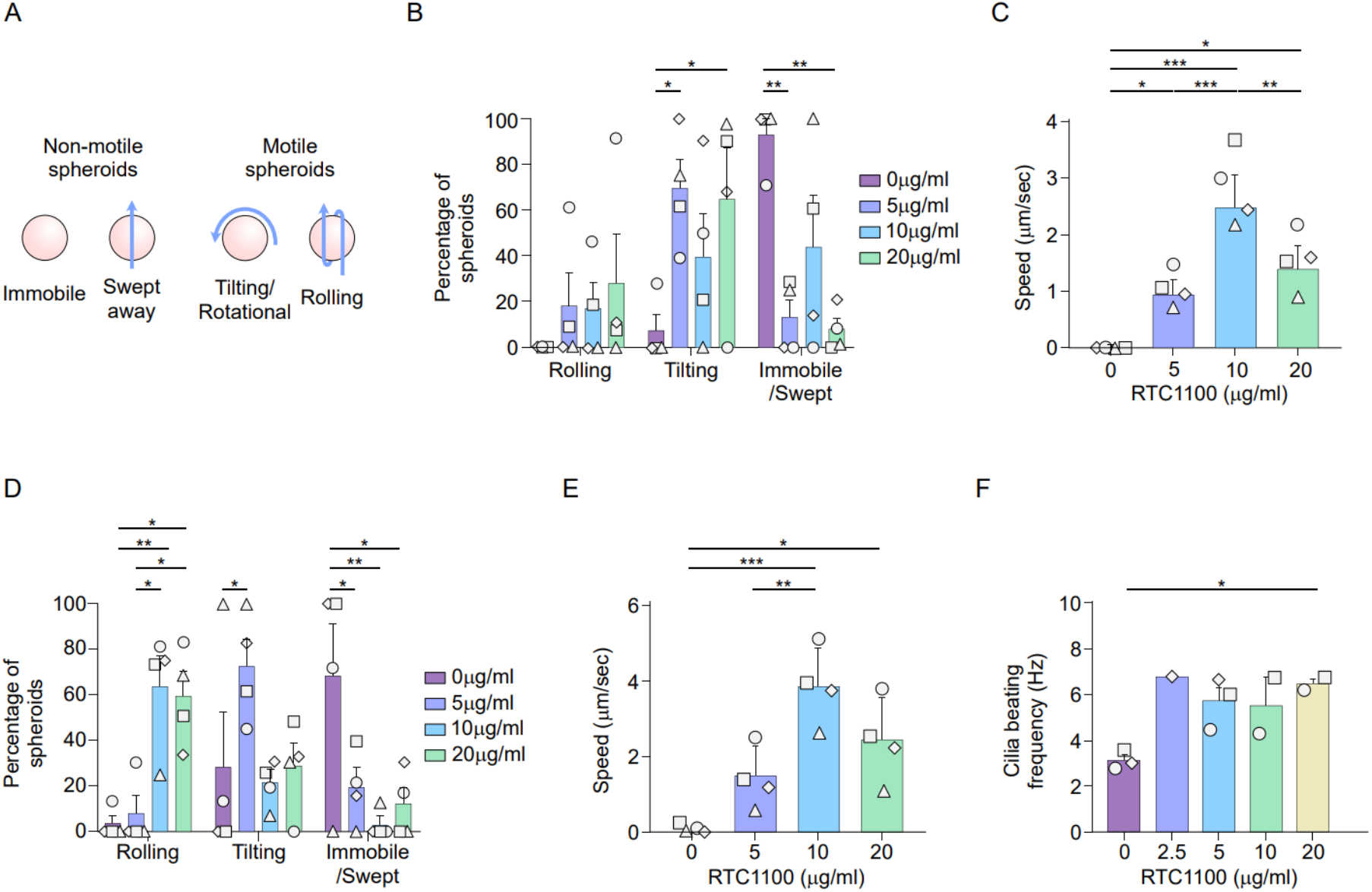
RTC1100 treatments improve spheroids motility and cilia beating frequency. A) Spheroids motility classification on this study. B-E) spheroids motility and rolling speed after the fourth dose (B and C) and sixth dose (D and E) of RTC1100 at 0, 5, 10 and 20 µg/ml. F) Cilia beating frequency after the sixth dose of RTC1100 at 0, 2.5. 5, 10 and 20 µg/ml. Each dot represents the average measurement of each patient.

To further quantify functionality, we measured translational speed of rolling spheroids over a 3-minute interval. At third dose, the rolling motion was just present in a small fraction of the spheroids (Figure 1B), with the higher speed in spheroids treated with 10 µg/mL (2.5µm/sec, Figure 1C). After six doses, untreated spheroids exhibited minimal movement (0.11 µm/s), whereas RTC-1100 increased mean speed to 1.5 µm/s, 3.9 µm/s and 2.5 µm/s at 5 µg/mL, 10 µg/mL and 20 µg/mL, respectively (Figure 1E). The greatest increase was observed at 10 µg/mL, consistent with peak effects on motility and coordination, indicating a dose-dependent effect with maximal response at 10 µm/mL.

### RCT1100 improves ciliary beat frequency in a dose-dependent manner

High-speed video microscopy was performed on all samples, except for one treated with 1.25 µg/mL RCT1100 due to insufficient spheroid quality for reliable analysis. At baseline, all samples demonstrated near-immotile cilia, with a mean (SD) CBF of 3.13 (0.723) Hz and a highly dyscoordinated ciliary beat pattern.

Following six doses of RCT1100 at concentrations of 2.5, 5, 10, and 20 µg/mL, CBF increased across all treatment groups, demonstrating a dose-dependent improvement (Figure 1). In addition to quantitative increases in CBF, qualitative assessment of HSVM recordings showed enhanced ciliary bending and improved synchrony of ciliary motion after treatment.

## Discussion

In this exploratory pilot study, we evaluated the effects of repeated *ex vivo* exposure to the investigational mRNA therapy RCT1100 in patient-derived nasal epithelial 3DE spheroids from individuals with PCD caused by genetically confirmed biallelic pathogenic variants in the DNAI1 gene. High-speed video microscopy demonstrated a dose-dependent increase in CBF, with values more than doubling in some samples. These findings were paralleled by improvements in coordinated spheroid movement observed by live-imaging, including increased rotational and rolling motion pattern, as well as enhanced translational speed, reaching up to approximately fourfold higher velocities compared with untreated controls.

*DNAI1* encodes a critical intermediate chain of the outer dynein arm, and biallelic pathogenic variants are classically associated with markedly reduced or absent ciliary motility^1^. Consistent with this phenotype, all baseline samples in the present study exhibited severely impaired ciliary function, reflected by low CBF and predominantly immotile spheroids. Following six doses of RCT1100, CBF increased across all evaluable concentrations, with relative improvements exceeding 100% in several samples. Although post-treatment CBF did not reach levels typically observed in healthy airway epithelium^5^, the magnitude of improvement is notable given the severity of the baseline dysfunction and the relatively short treatment duration.

Importantly, increases in CBF were accompanied by qualitative and functional improvements in coordinated ciliary activity. Untreated spheroids largely remained static, whereas treated spheroids exhibited pronounced rotational and rolling motion patterns demonstrated in our previous study^4^ indicative of improved ciliary synchrony and effective force generation. The free-field configuration of the 3DE system enables simultaneous assessment of both rotational and translational motion without mechanical constraints, providing a more integrated functional readout than CBF alone. The observed increases in translational velocity further support the conclusion that RCT1100 enhances overall ciliary performance.

A dose–response relationship was observed, with higher RCT1100 concentrations generally associated with more sustained improvements following repeated dosing. While the 5 µg/mL concentration elicited early gains, these were not maintained at the final time point, in contrast to the more consistent responses observed at 10 and 20 µg/mL. This pattern may reflect differences in mRNA uptake, intracellular persistence, or turnover of the DNAI1 protein, and underscores the importance of dose optimization and repeat administration for mRNA-based therapies targeting motile cilia. Inter-individual variability in treatment response was also noted, potentially attributable to differences in genotype, epithelial cell integrity, or spheroid formation efficiency.

This study also highlights the utility of 3DE spheroids as a rapid and sensitive way for evaluating therapeutic responses in PCD. Compared with conventional air-liquid interface cultures based on cell differentiation^4^, spheroids in the present study are formed directly from undifferentiated cells derived from nasal brushings and require minimal manipulation and are suitable for functional analysis within few days. The concordance between improvements in CBF and enhanced spheroid motility supports the validity of this model as a functional assay for preclinical and early-phase translational studies.

Several limitations warrant consideration. The sample size was small, and a proportion of samples were excluded due to contamination or inadequate viability, reflecting the technical challenges associated with primary airway epithelial cultures. The study was descriptive in nature and did not include formal statistical testing, limiting the strength of inference. In addition, while restoration of ciliary motility is a key prerequisite for effective mucociliary clearance, downstream functional outcomes such as mucus transport and durability of response were not assessed.

Despite these limitations, this study provides proof-of-concept evidence that mRNA-based replacement of DNAI1 can restore key aspects of ciliary function in patient-derived airway epithelium. Together with prior preclinical findings demonstrating rescue of ciliary activity in air–liquid interface models^3^ these results support continued clinical development of RCT1100 and further exploration of mRNA therapeutics for genetically defined subtypes of PCD. Future studies incorporating larger cohorts, standardized dosing regimens, and additional functional endpoints at various predefined timepoints will be essential to establish the therapeutic potential and clinical relevance of this approach.

## Abbreviations

3DE: Three-Dimensional Explant
CBF: ciliary beat frequency
CBP: ciliary beat pattern
DNAI1: Dynein Axonemal Intermediate Chain 1
LNP: lipid nano particle
MCC: Mucociliary clearance
PCD: Primary ciliary dyskinesia
ppFEV1: percent predicted Forced Expiratory Volume in the first second
SD: Standard deviation
SORT: selective organ targeting
TEM: Transmission electron microscopy

## Declarations

### Ethics and consent to participate

The clinical trial was approved and registered at EUCT no.: 2023-510521-14-00. All patients have signed an informed consent form

### Consent for publication

All patients have given written consent for the data and videos to be published.

### Availability of data and materials

Data and materials are available upon reasonable request.

### Authors contributions

Patient recruitment and enrollment: TQ, CMTN

Sample Collection: CMTN, JKM

Spheroid cultivation and dosing: CHC, LN, CMTN, JKM

Data collection: CMTN, CHC, LN, SGJ, JKM, TQ

Data analysis: CMTN, CHC, JKM, STC

Research project management: STC, KGN, JKM, CMTN, CHC

Writing original draft: CMTN, CHC, JKM, KGN, STC

All authors read and approved the final manuscript

